# OncoVarAgent: An Autonomous Research Agent for Discovering Potential Therapeutic Strategies for Somatic Cancer Variants Lacking Established Guidance

**DOI:** 10.64898/2026.11.24.769026

**Authors:** Zelin Wang, Minfang Cai, Ziling Zhan, Yidian Dong, Zhenyu Lin, Zhongyan Chen, Yuanyuan Yin, Yiqing Zheng

**Affiliations:** Department of Otolaryngology, Sun Yat-Sen Memorial Hospital, Sun Yat-Sen University, Guangzhou, China; Guangdong Provincial Key Laboratory of Cancer Pathogenesis and Precision Diagnosis and Treatment, Joint Big Data Laboratory, Department of Medical Oncology, Shenshan Medical Center, Sun Yat-sen Memorial Hospital, Sun Yat-sen University, Shanwei, China; Guangdong Provincial Key Laboratory of Malignant Tumor Epigenetics and Gene Regulation, Guangdong-Hong Kong Joint Laboratory for RNA Medicine, Medical Research Center, Sun Yat-sen Memorial Hospital, Sun Yat-sen University, Guangzhou, China; Department of Precision Medicine, Shenshan Medical Center, Sun Yat-sen Memorial Hospital, Sun Yat-sen University, Shanwei, China

**Keywords:** Precision oncology, Somatic variants, Variant interpretation, AI agent

## Abstract

**Background:** Variant interpretation remains a major challenge in precision oncology, particularly for somatic cancer variants that lack established standard-of-care therapeutic guidance. However, relevant evidence is often dispersed across biomedical literature and clinical trial databases, making manual identification and synthesis labor-intensive and difficult to scale. This study aims to develop an efficient LLM-based agent to automatically identify and synthesize evidence to support somatic cancer variant interpretation in precision oncology.

**Methods:** We developed OncoVarAgent, powered by GPT-5.4, an autonomous artificial intelligence agent designed to discover potential therapeutic strategies for somatic cancer variants without standard-of-care drugs. OncoVarAgent uses the ReAct reasoning-and-acting framework to dynamically query PubMed and ClinicalTrials.gov, analyze heterogeneous evidence, and generate structured reports. We evaluated OncoVarAgent using a benchmark cohort of 30 AMP Tier II–III somatic cancer variants across nine tumor types. Generated reports were independently reviewed by domain experts across predefined dimensions, including scientific accuracy, evidence relevance, therapeutic hypothesis plausibility, clinical utility, and completeness.

**Results:** Among the 30 evaluated variants, OncoVarAgent identified candidate therapeutic evidence for 19 variants. Across these 19 variants, it reported a median of 9 drugs per variant, supported by a median of 7 publications and 4 clinical trials. The median runtime was 2.46 minutes and the median total cost was $0.72 per variant. Independent expert review showed generally favorable evaluations across five predefined dimensions, with mean scores ranging from 3.68 to 4.11 on a five-point scale and statistically significant positive inter-expert concordance across all dimensions.

**Conclusions:** OncoVarAgent demonstrates the feasibility of using an autonomous AI agent for structured therapeutic evidence retrieval and synthesis to assist somatic variant interpretation in precision oncology.

## Introduction

The advent of cancer genomic profiling has fundamentally reshaped the landscape of precision oncology, enabling the systematic identification of somatic alterations across hundreds of cancer-related genes^1,2^. This technology holds the promise of tailoring treatments to the specific molecular drivers of a patient’s tumor, a paradigm that has led to notable successes for a subset of patients with well-characterized mutations such as EGFR L858R and BRAF V600E^3,4^. However, a major hurdle to the broader adoption of precision oncology is the difficulty in variant interpretation. Although several knowledge bases, including OncoKB^5^, CIViC^6^, PMKB^7^ and CGI^8^, have been developed to support clinicians, they primarily focus on annotating FDA-approved drugs. For example, OncoKB has annotated only 159 drugs for over 10,000 alterations, leaving the vast majority of variants without potential therapeutic clues. This is particularly problematic for Molecular Tumor Boards (MTBs), which are often confronted with therapeutically ambiguous variants. In such cases, MTBs must perform extensive literature reviews to find evidence that could be decisive for recommending an off-label therapy or clinical trial enrollment for advanced-stage patients who have exhausted standard treatment options.

The rapid advancements in large language models (LLMs) and agent-based AI systems present a transformative opportunity for medicine^9-13^. These technologies have been demonstrated on many varieties of clinical tasks such as patient-to-clinical trial matching^14^, clinical text summarization^15^ and clinical decision-making^16^. More recently, a RAG-LLM framework leveraging the MOAlmanac database has been developed to provide accurate FDA-approved therapy recommendations^17^. Similarly, RAG-based LLMs like OpenEvidence have also emerged to assist with medical literature searches and clinical decision support. Despite these advancements, there is a lack of AI tools available to assist MTBs in automating the exhaustive task of literature reviews and clinical trial matching. This manual process is incredibly time-consuming, typically requiring several days to weeks, and severely delays or prevents the discovery of potential therapeutic strategies for advanced cancer patients.

Here, we present OncoVarAgent, an autonomous AI agent designed to systematically identify potential therapy strategies for variants lacking established guidelines. By deploying a reasoning-driven agent to dynamically query, analyze, and synthesize information from PubMed and ClinicalTrials.gov based on ReAct (Reasoning and Acting) framework, OncoVarAgent automates therapeutic evidence retrieval and synthesis for cancer variant interpretation within minutes. We demonstrate that this agent, powered by GPT-5.4, may be useful for clinicians who need to query, review and synthesize large-scale biomedical literature and clinical trial information for cancer variant interpretation. This was evaluated in a benchmark cohort of 30 AMP Tier II–III somatic cancer variants across nine tumor types. OncoVarAgent provides a structured and scalable approach augment evidence retrieval and synthesis for variants lacking established guidance in precision oncology.

## Methods

### The OncoVarAgent Framework

We developed OncoVarAgent, an autonomous agent designed to systematically identify potential therapy strategies for variants lacking standard-of-care drugs. The entire workflow is implemented in Python using the LangGraph library to construct a stateful computational graph. The framework is orchestrated as a series of interconnected nodes, each performing a specific task, from initial data annotation to final evidence synthesis. GPT-5.4 is used in this study with temperature as 0. The overall architecture is depicted in Fig. 1.

**Fig. 1.**
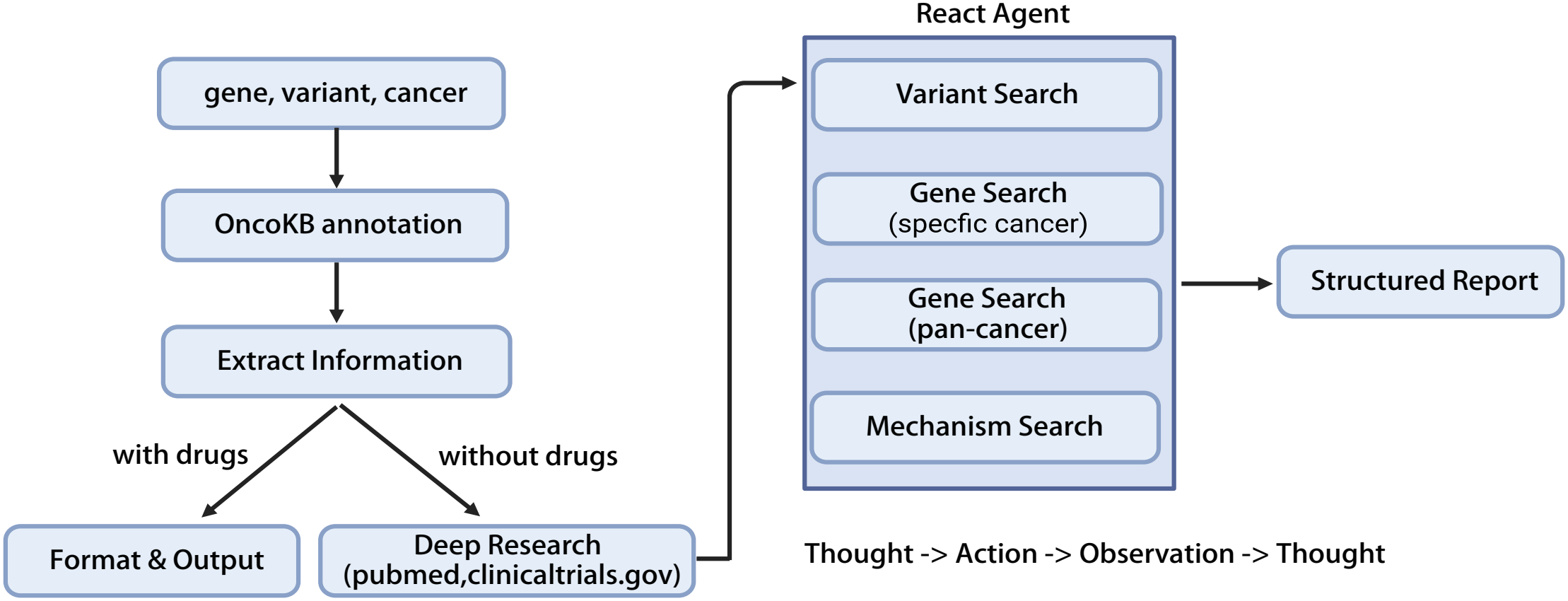
The workflow of the OncoVarAgent framework. The process initiates with variant annotation via OncoKB. If a variant lack established drug associations, it triggers the Deep Research module, which is powered by a ReAct agent. This agent systematically queries external databases like PubMed and ClinicalTrials.gov through four-staged search strategies. The agent operates on an iterative ‘Thought -> Action -> Observation’ cycle to synthesize evidence, ultimately generating a structured evidence report.

### Input Data and Baseline Annotation

The workflow initiates with a standard input file, containing essential columns for gene symbol (e.g., ‘Hugo_Symbol’), protein change (e.g., ‘HGVSp_Short’), and cancer type. To establish a baseline of clinical evidence, this file is first processed by the MafAnnotator.py from OncoKB. The annotator enriches the input variants with structured information from the OncoKB, including oncogenicity status, mutation effect, and therapeutic implications (AMP/ASCO/CAP^18^ Tier). When a variant is annotated as ‘Oncogenic’, ‘Likely Oncogenic’, or has an unknown oncogenic status, and critically, has no pre-existing drug associations in OncoKB, it is prioritized for in-depth analysis by the deep research agent.

### The Deep Research Agent

We implemented a specialized evidence-discovery agent based on the ReAct framework. The agent was designed to emulate a structured oncology literature-review workflow for somatic cancer variants that lack established therapeutic guidance. It was powered by GPT-5.4 and equipped with two custom tools: pubmed_search, which programmatically queries the NCBI PubMed database and returns up to 20 abstracts per query, and query_clinical_trials, which queries the ClinicalTrials.gov API for active interventional studies with recruitment statuses including RECRUITING, NOT_YET_RECRUITING, ACTIVE_NOT_RECRUITING, and ENROLLING_BY_INVITATION, returning up to 20 records per request.

The agent followed a predefined four-phase workflow. Baseline functional annotation from OncoKB, including mutation effect and mutation-effect description, was provided to the agent before evidence exploration. In Phase 1, the agent performed variant-level functional characterization and searched for direct therapeutic evidence for the specific variant. This phase included two required PubMed searches: one restricted to the patient’s cancer type and one broader pan-cancer search. After reviewing both searches together with the OncoKB annotation, the agent was required to explicitly determine whether the variant was best interpreted as gain-of-function, loss-of-function, or functionally uncertain. If the variant function remained uncertain, the agent terminated the therapeutic exploration to avoid unsupported therapeutic inference.

In Phase 2, the agent performed a gene-focused therapeutic search within the patient’s cancer type using therapy-oriented PubMed queries. The goal of this phase was to identify drugs, drug classes, downstream pathways, or therapeutic vulnerabilities associated with the altered gene in the same disease context. In Phase 3, the agent broadened the search to a pan-cancer context to identify therapeutic evidence or approvals that may be relevant across tumor types. In Phase 4, the agent performed a mechanistic deep dive. Unlike the earlier phases, which used predefined gene-, variant-, and cancer-type queries, Phase 4 used biological mechanisms inferred from the preceding phases and the gain- or loss-of-function interpretation to construct mechanism-focused follow-up searches. These searches were intended to identify indirect therapeutic strategies, pathway dependencies, synthetic-lethality relationships, resistance or sensitivity mechanisms, and evidence from other cancer contexts that could support biologically plausible therapeutic hypotheses.

Clinical trial search was implemented as a reactive validation step rather than a standalone screening step. Whenever a PubMed search identified a potential drug, drug class, pathway inhibitor, or targetable biological mechanism, the agent was required to immediately call query_clinical_trials using structured parameters such as intervention and condition. If a specific clinical trial query returned no relevant results, the agent was instructed not to stop, but to broaden the trial search by relaxing the query, for example by searching only the intervention or therapeutic class. This design was intended to link literature-derived therapeutic hypotheses to the current interventional trial landscape while reducing untargeted trial retrieval.

The agent was constrained to single-tool execution per reasoning step. Before each tool call, it was required to summarize the result of the preceding action, including relevant PMID or NCT identifiers when available, and then state the rationale for the next action. This enforced an iterative Thought–Action–Observation cycle and provided an auditable reasoning trace for downstream review. PubMed queries involving Boolean OR terms were required to use explicit parentheses, and all PubMed and ClinicalTrials.gov calls were capped at 20 returned records per request.

At completion, the agent synthesized the retrieved evidence into a structured mini-review with predefined sections: Executive Summary, Evidence Synthesis, Conclusion, and Curated Evidence Lists. Findings in the Evidence Synthesis section were grouped by therapeutic strategy or drug class rather than by search order. For each therapeutic hypothesis, the agent was required to state the biological rationale, summarize the supporting evidence, distinguish the level of evidence, and cite relevant PubMed IDs and ClinicalTrials.gov identifiers inline. The final curated PMID and NCT lists were required to exactly match the identifiers cited in the Evidence Synthesis section, thereby enabling transparent evidence tracing. The full prompt was provided in Supplementary Methods.

### Report Generation Agent

Upon completion of the deep research, the Deep Research Agent produces a textual evidence summary and a curated list of supporting literature and clinical trials. These outputs are passed to the final report generation agent. This agent’s role is strictly defined as a non-judgmental data processor. Its prompt explicitly forbids it from making new scientific inferences and instead directs it to perform a series of deterministic formatting and extraction tasks. Specifically, the prompt instructs the model to: (1) reformat the researcher’s free-text summary into a standardized Markdown structure with predefined Summary, Results, and Conclusion sections; (2) generate a new, concise two-to-three sentence brief report for high-level review; and (3) perform structured entity extraction to populate fields for drugs, PubMed IDs, and Clinical Trial IDs in specific string formats. To guarantee adherence to the data model, the prompt concludes by providing the exact JSON schema to guide the final output. Clinical trial number: not applicable.

### Benchmark Dataset Construction

We constructed a benchmark dataset from somatic cancer variants curated at Department of Precision Medicine, Shenshan Medical Center, Sun Yat-sen Memorial Hospital, Sun Yat-sen University. Candidate variants were derived from molecular profiling reports of patients with solid tumors and were reviewed. We focused on AMP Tier II and Tier III variants because these alterations are frequently lacking straightforward standard-of-care therapeutic guidance. From the eligible Tier II–III variant pool, 30 somatic variants were randomly selected.

### Expert Evaluation of OncoVarAgent Reports

We designed a five-dimensional scoring rubric to evaluate the quality of OncoVarAgent-generated reports. The five dimensions were: scientific accuracy, evidence relevance, therapeutic hypothesis plausibility, clinical utility and completeness. Each dimension was scored on a five-point Likert scale, with 1 indicating poor performance and 5 indicating excellent performance. Scientific accuracy assessed whether the report correctly described the biological function of the variant. Evidence relevance assessed whether the cited PubMed articles and ClinicalTrials.gov records were directly relevant to the variant. Therapeutic hypothesis plausibility assessed whether proposed strategies were biologically plausible. Clinical utility assessed whether the report could provide useful information for variant interpretation. Completeness assessed whether the report captured the major therapeutic directions, relevant mechanisms, and ongoing clinical trials available for the variant or its implicated pathway. For each dimension, mean scores and 95% confidence intervals were calculated for each expert. Concordance between the two experts was assessed using Pearson correlation coefficients across the evaluated variants. The full scoring rubric, including the detailed criteria for scores from 1 to 5 in each dimension, was provided in Supplementary Methods.

### Implementation

OncoVarAgent was implemented as a modular system supporting interactive web-based use, local offline execution, and batch analysis. The web interface was developed using React and Tailwind CSS, providing a responsive front end for variant submission and report review. The backend service was primarily built with FastAPI, which handled task execution. To facilitate local use without deploying the full web stack, we also developed an independent offline application using Streamlit, enabling users to run OncoVarAgent locally through a lightweight graphical interface. In addition, a standalone Python script was provided for batch processing of multiple variants from the command line. Further details were available at https://github.com/charlin90/OncoVarAgent.

## Results

### OncoVarAgent workflow

OncoVarAgent was implemented as a graph-based, LLM-powered framework that integrates variant annotation, evidence retrieval, agentic reasoning, and structured report synthesis (Fig. 1). For each input variant, the system first performs OncoKB-based annotation and then applies a routing strategy. Variants with existing OncoKB-supported therapeutic associations or likely neutral classifications are formatted directly into the final report, whereas variants lacking direct therapeutic annotation are routed to a ReAct-style deep research module. This module iteratively queries PubMed and ClinicalTrials.gov to retrieve literature and clinical trial evidence. The final synthesizer integrates baseline annotations with retrieved evidence to generate a structured report containing therapeutic hypotheses, mechanistic rationale, supporting PubMed identifiers, and relevant clinical trial identifiers.

### Overview of the 30-variant benchmark cohort

To evaluate the performance of OncoVarAgent, we curated a benchmark dataset comprising 30 cancer variants across nine tumor types (Supplementary Table 1). Non-small cell lung cancer represented the largest subgroup (14/30, 46.7%), followed by breast cancer (4/30, 13.3%) and gastric cancer (3/30, 10.0%) (Fig. 2A). Most variants were classified as AMP Tier II (24/30, 80.0%), with the remaining variants classified as Tier III (6/30, 20.0%) (Fig. 2B). The dataset included nonsense (12/30, 40.0%), missense (9/30, 30.0%), and frameshift alterations (9/30, 30.0%) (Fig. 2C). The cohort covered a broad gene spectrum, with TP53 being the most frequent gene (3/30, 10.0%), followed by ARID2 and BMPR1A (2/30 each, 6.7%); the remaining genes were represented by one variant each (Fig. 2D).

**Fig. 2.**
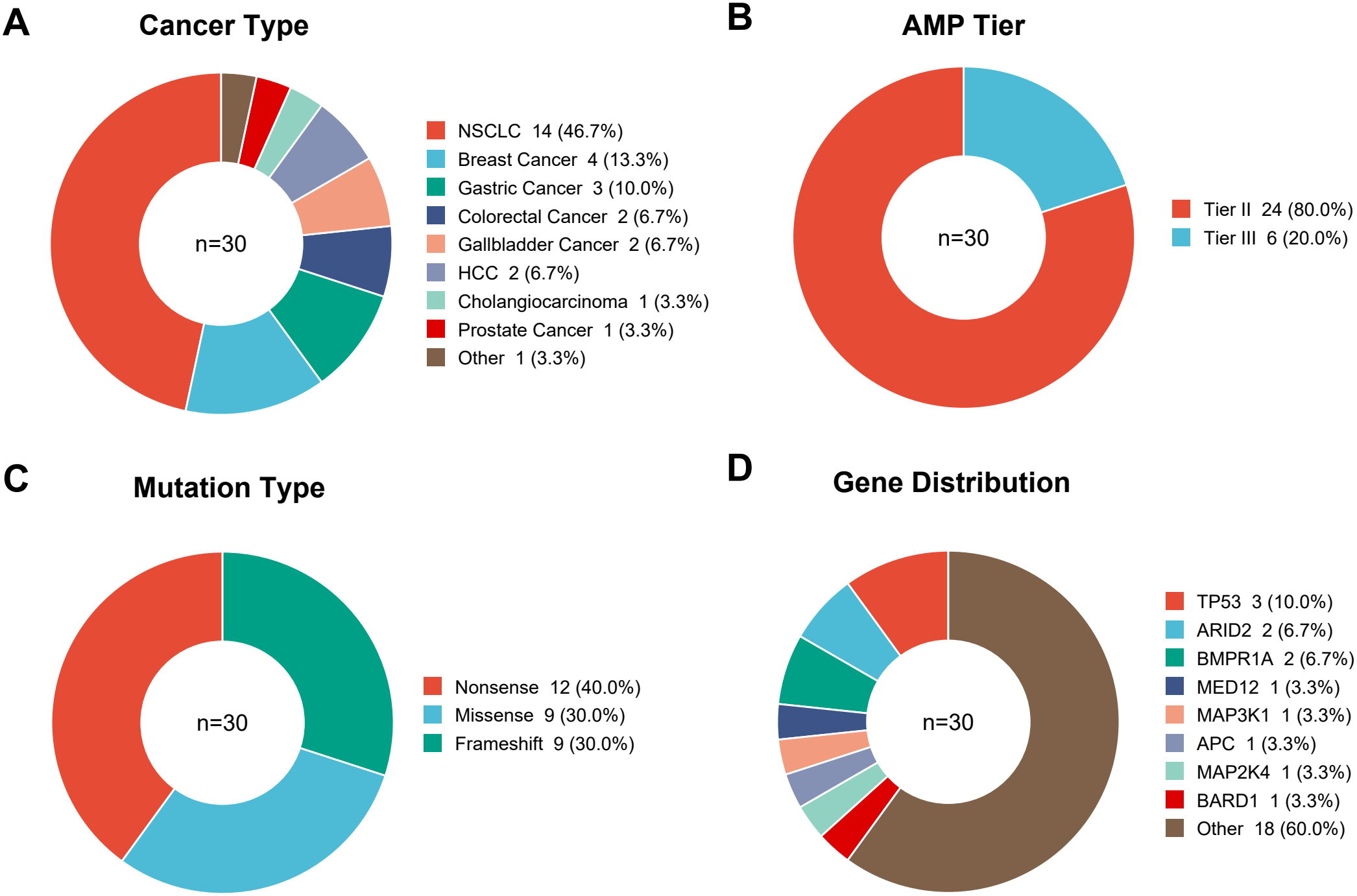
Composition of the benchmark dataset. The benchmark dataset comprised 30 somatic cancer variants across nine tumor types. Donut plots show the distribution of variants by cancer type (A), AMP tier (B), mutation type (C), and altered gene (D).

### Therapeutic evidence retrieval, synthesis, and computational burden

Among the 30 variants, 5 (16.7%) had drug evidence identified during the initial OncoKB annotation step and were therefore not subjected to the deep research module. Of the remaining 25 variants evaluated by deep research, OncoVarAgent identified potential therapeutic hypotheses for 19 variants (63.3% of the full cohort), whereas 6 variants (20.0%) without effective supporting evidence meeting the predefined reporting criteria was identified (Fig. 3A). Across the 19 variants with candidate therapeutic evidence, OncoVarAgent reported a median of 9 drugs per variant, supported by a median of 7 curated publications and 4 curated clinical trial identifiers (Fig. 3B). The computational burden was modest, with a median runtime of 2.46 minutes per variant and a median total cost of $0.72 per variant (Fig. 3C).

**Fig. 3.**
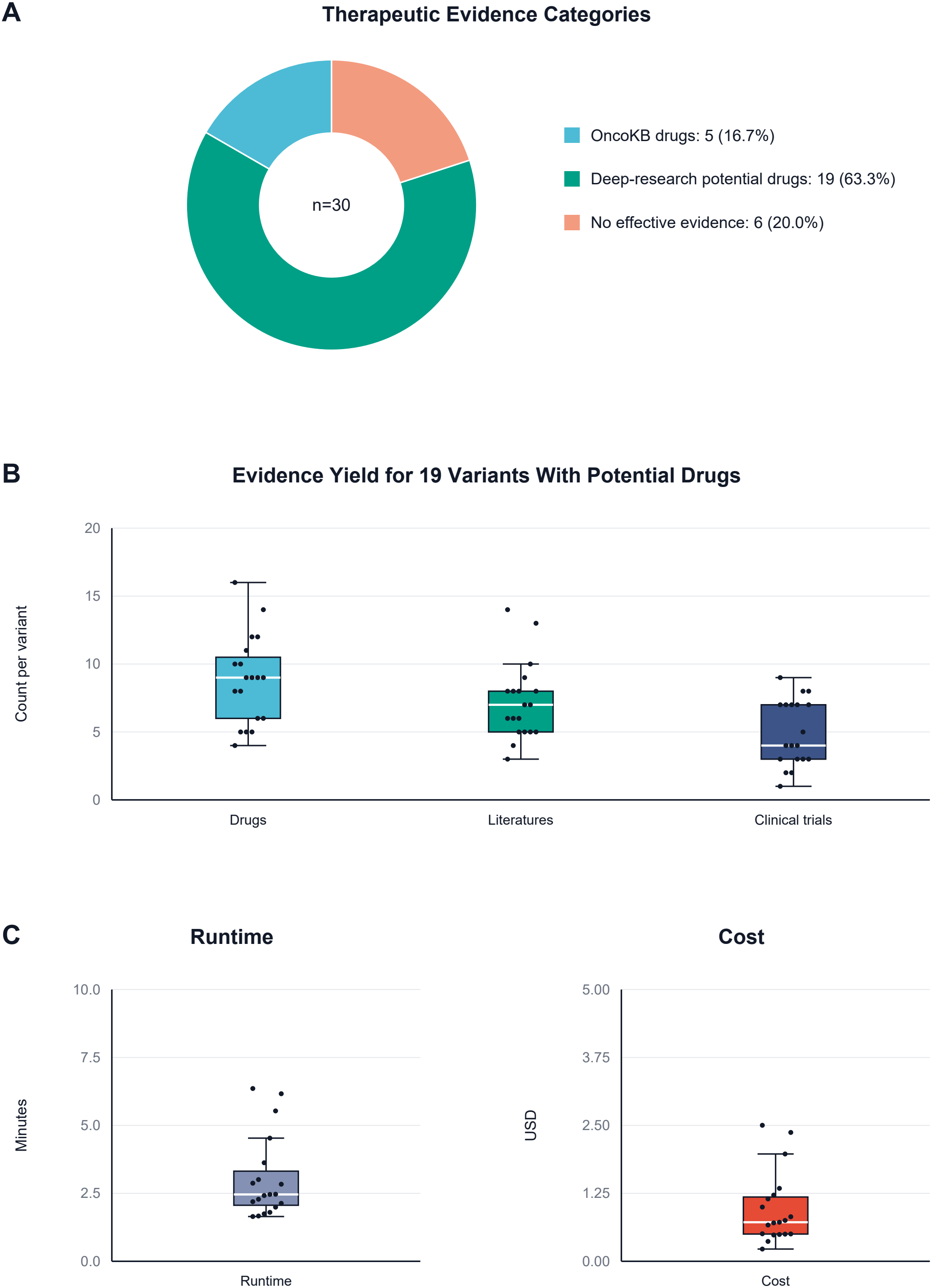
Therapeutic evidence retrieval and computational burden of OncoVarAgent. (A) Distribution of therapeutic evidence categories across the 30-variant benchmark cohort. (B) Evidence yield among the 19 variants with reportable therapeutic evidence, including the number of candidate drugs, curated supporting publications, and curated clinical trial identifiers per variant. (C) Runtime and computational cost across the 19 variants.

### Expert assessment of OncoVarAgent reports

Expert review was performed for the 19 variants for which OncoVarAgent identified potential therapeutic evidence. Across the five predefined evaluation dimensions, both experts assigned generally high scores, with mean scores ranging from 3.68 to 4.11 on a five-point scale (Fig. 4A, Supplementary Table 2 and 3). The highest mean scores were observed for scientific accuracy, whereas relatively lower scores were observed for evidence relevance and clinical utility. Score concordance between the two experts was positive across all dimensions, with Pearson correlation coefficients of 0.84 for scientific accuracy, 0.69 for evidence relevance, 0.76 for therapeutic hypothesis plausibility, 0.92 for clinical utility, and 0.73 for completeness; all correlations were statistically significant (p < 0.001, Fig. 4B-F). These results suggest that the agent-generated reports were generally rated favorably among cases with reportable therapeutic evidence.

**Fig. 4.**
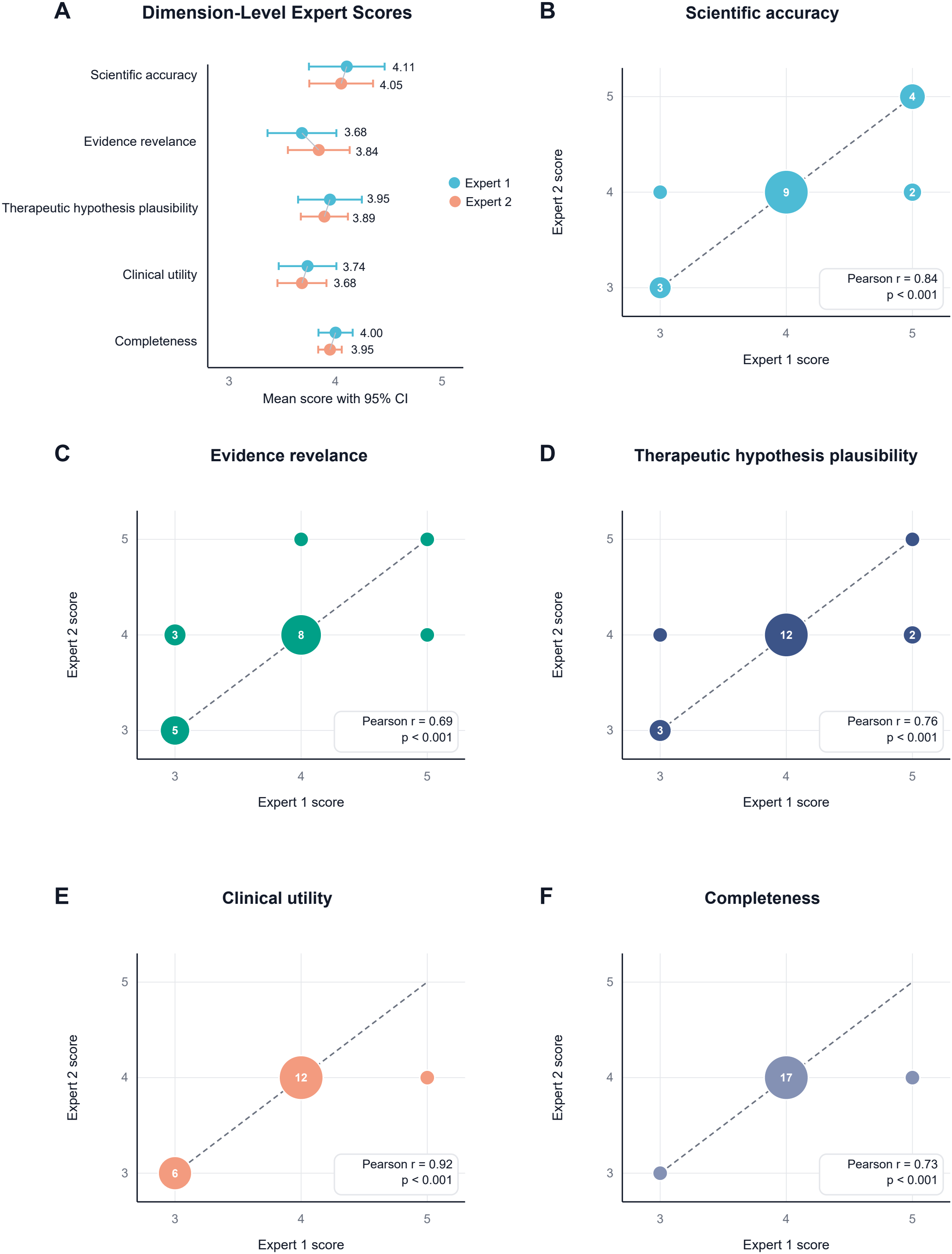
Expert assessment of OncoVarAgent-generated evidence reports. (A) Mean expert scores across five predefined evaluation dimensions for the 19 variants with reportable therapeutic evidence. Points indicate mean scores, and error bars indicate 95% confidence intervals. (B–F) Concordance between the two independent experts for scientific accuracy (B), evidence relevance (C), therapeutic hypothesis plausibility (D), clinical utility (E), and completeness (F). Each point represents an expert-score pair for one variant; larger bubbles indicate multiple variants with the same score combination. Dashed lines indicate perfect concordance between the two experts. Pearson correlation coefficients and corresponding p values are shown in each panel.

### Case-level interpretation of an RBM10 loss-of-function variant

We present the case of the RBM10 N446Kfs*35 variant in Non-Small Cell Lung Cancer (NSCLC), a ‘Likely Oncogenic’ mutation with loss of function for which the OncoKB provides no standard variant-specific therapy. OncoVarAgent proposed a potential benefit from immune checkpoint inhibitors in *EGFR*-wildtype tumors, citing data associating *RBM10* loss-of-function with immunogenic features^19^, while simultaneously flagging the same mutation as a marker of potential resistance to osimertinib in *EGFR*-mutant contexts based on multiple retrospective studies^20,21^. importantly, it pinpointed the RAC1B pathway, negatively regulated by RBM10, as the underlying resistance mechanism and a potential combination target^22^. Furthermore, the agent identifies novel vulnerabilities, such as a potential sensitivity to PRMT5 pathway inhibitors targeting spliceosome dependencies^23^, and a synthetic lethal relationship with WEE1 inhibitors in *RBM10*-deficient models^24^. Most actionably, it connected a preclinical rationale for targeting the S1P pathway to prevent brain metastasis^25^ with a recruiting Phase 2 clinical trial for the S1P modulator fingolimod (NCT06424067). Additional details were provided in Supplementary File 1.

## Discussion

In this study, we developed and demonstrated OncoVarAgent, an autonomous AI agent designed to address the critical variant interpretation bottleneck in precision oncology. Our work moves beyond traditional, static knowledge bases by creating a dynamic, reasoning-driven agent that actively discovers and synthesizes therapeutic evidence for somatic variants lacking established clinical guidance.

Unlike systems designed to confirm on-label therapies^17^, OncoVarAgent is specifically engineered for the frontier of clinical genomics—where evidence is nascent, indirect, or buried within disparate preclinical and clinical data. The RBM10 N446Kfs*35 case study is a powerful illustration of this capability. OncoVarAgent synthesized a loss-of-function tumor-suppressor alteration into a set of context-dependent hypotheses, including attenuated EGFR-TKI benefit^20,21^, possible immune responsiveness in EGFR-wildtype tumors^19^, and investigational vulnerabilities involving RAC1B^22^, PRMT5^23^, WEE1^24^, and S1P signaling^25^. The agent appropriately framed them as evidence-linked, mechanism-based hypotheses for expert review, trial consideration, and future validation.

The potential clinical utility of OncoVarAgent is promising. For MTBs, which are frequently tasked with interpreting complex genomic reports under time constraints, the agent offers a means to systematically and rapidly generate a comprehensive evidence landscape. These outputs should be viewed not as definitive treatment recommendations, but as prioritized hypotheses to assist somatic variant interpretation in precision oncology. By automating the laborious process of searching PubMed and ClinicalTrials.gov and structuring the output, OncoVarAgent may reduce the manual evidence-query burden and facilitate pre-MTB case preparation.

Despite these promising capabilities, it is important to acknowledge the inherent limitations of the current framework. The agent’s performance is fundamentally dependent on the quality and accessibility of the underlying data sources. In the current implementation, literature retrieval is based on PubMed abstracts rather than full-text articles; therefore, relevant evidence described only in full-text sections, supplementary materials, figures, or non-indexed sources may be missed. Furthermore, like all LLM-based systems, OncoVarAgent is susceptible to “hallucinations” or misinterpretations. Therefore, OncoVarAgent should be viewed as a decision-support tool designed to assist clinicians in identifying, retrieving, and synthesizing relevant evidence from large-scale biomedical literature and clinical trial databases. Final interpretation and clinical decision-making must always remain the responsibility of human experts.

Future work will focus on several key areas to enhance the agent’s robustness and utility. First, expanding the suite of tools to include additional data sources, such as genomic databases (e.g., cBioPortal^26^, TCGA^27^), drug databases^28^, and conference proceedings, could provide a more holistic view of the evidence landscape. Second, developing more sophisticated methods for the agent to automatically assess and rank the level of evidence would further refine the actionability of its reports. Finally, integrating variant effect prediction tools such as AlphaMissense^29^ and AlphaGenome^30^ may help improve functional interpretation for variants with limited experimental or clinical evidence.

In conclusion, OncoVarAgent demonstrates the potential of agentic AI systems to move beyond static information retrieval toward structured evidence retrieval and synthesis in clinical cancer genomics. By systematically navigating and synthesizing heterogeneous biomedical data, it provides a scalable framework to assist cancer variant interpretation.

## Declarations

### Author contributions

Y.Z. conceived and supervised the project and acquired funding. Z.W. designed and developed the software, performed the analyses, and wrote the original draft. M.C., Z.Z, Y.D, Z.L., Z.C., and Y.Y. contributed to the study design and validation. All authors contributed to the review and editing of the manuscript and approved the final version.

### Ethics approval and consent to participate

The benchmark variants used in this study were retrospectively curated from deidentified somatic variant information at the Department of Precision Medicine, Shenshan Medical Center, Sun Yat-sen Memorial Hospital, Sun Yat-sen University. This retrospective study was reviewed and approved by the Medical Ethics Committee of Shenshan Medical Center, Sun Yat-sen Memorial Hospital, Sun Yat-sen University. The requirement for informed consent to participate was waived by the Medical Ethics Committee of Shenshan Medical Center, Sun Yat-sen Memorial Hospital, Sun Yat-sen University. s’s

### Consent for publication

Not applicable.

### Availability of data and materials

The OncoVarAgent web application is publicly available at http://oncovaragent.databio1.com. The offline Streamlit application and command-line scripts for batch execution are available at https://github.com/charlin90/OncoVarAgent. Additional data generated or analyzed during the current study are available in Supplementary Materials.

## Acknowledgements

Not applicable.

## Supplementary Tables

**Supplementary Table 1**. Benchmark cohort characteristics and OncoVarAgent output summary.

**Supplementary Table 2**. Expert 1 evaluation scores for OncoVarAgent-generated reports.

**Supplementary Table 3**. Expert 2 evaluation scores for OncoVarAgent-generated reports.

