## Supplementary material for "OncoVarAgent: An Autonomous Research Agent for Discovering Potential Therapeutic Strategies for Somatic Cancer Variants Lacking Established Guidance": Supplementary File 1.pdf

### 1. Executive Summary:

RBM10 N446Kfs*35 in NSCLC is best interpreted as a **loss-of-function (LoF)** truncating variant. No direct variant-specific therapeutic reports were found, but RBM10-deficient NSCLC/LUAD has emerging **mechanism-based vulnerabilities**: strongest preclinical evidence supports **WEE1 inhibition**, **S1P pathway inhibition/fingolimod** for brain-metastatic biology, and **splicing/RAC1B-directed strategies**; RBM10 alterations also appear associated with **weaker EGFR-TKI benefit** and context-dependent **immunotherapy signals** (PMID: 39080280, 40069781, 40548642, 38323283, 38912481, 39138107, NCT: NCT06424067).

### 2. Evidence Synthesis:

#### A. Exploiting replication stress with WEE1 inhibition

**Therapeutic hypothesis:** Target RBM10-deficient LUAD/NSCLC with **WEE1 inhibitors**.
**Rationale and evidence:** A recent mechanistic LUAD study identified **synthetic lethality between RBM10 loss and WEE1**, showing that RBM10-deficient cells have replication-fork/replication-stress vulnerabilities and are sensitized to WEE1 inhibition both **in vitro and in vivo**. This is the most directly RBM10-LoF-driven therapeutic strategy identified in lung adenocarcinoma (PMID: 39080280).
**Strength:** Strong **preclinical** evidence in RBM10-deficient LUAD.
**Clinical translation status:** In this search workflow, I did not identify an active NSCLC-specific adavosertib trial tied to RBM10 status, so this remains a compelling but **investigational hypothesis without variant-specific clinical validation** in NSCLC from the retrieved trial set.

#### B. Targeting sphingolipid/S1P signaling, especially in EGFR-mutant LUAD brain-metastatic settings

**Therapeutic hypothesis:** Use **S1P-pathway inhibition**, particularly **fingolimod**, in RBM10-deficient EGFR-mutant LUAD with metastatic/brain-metastatic biology.
**Rationale and evidence:** RBM10 deficiency was shown to alter **GALC splicing**, increase **sphingosine-1-phosphate (S1P)** production, enhance BBB permeability, and promote brain metastasis. In that model, **fingolimod** and **RU-SKI-43** suppressed brain-metastatic growth, directly linking RBM10 loss to a druggable pathway (PMID: 40069781).
**Strength:** Strong **preclinical** evidence.
**Clinical translation status:** There is an active phase II lung cancer trial of **fingolimod** in patients with NSCLC/SCLC after chemo-immunotherapy, providing a real clinical avenue for this pathway even though the trial is not biomarker-selected for RBM10 (NCT: NCT06424067). No active interventional trial was found for RU-SKI-43 in the performed searches.

#### C. Counteracting RBM10-loss-driven splicing dysregulation: RAC1B axis and PRMT5-pathway strategies

**Therapeutic hypothesis 1:** Target **RAC1B/splicing-dependent oncogenic output** to enhance EGFR inhibitor efficacy.
**Rationale and evidence:** RBM10 was identified as a **negative regulator of RAC1B** in EGFR-mutant LUAD. RAC1B promoted tumor growth and stabilized EGFR, and **targeting RAC1B suppressed tumor growth and enhanced EGFR inhibitor efficacy** in organoid/xenograft models. This makes RAC1B a biologically coherent downstream target for RBM10-LoF tumors,

particularly in EGFR-mutant disease (PMID: 40548642).
**Strength: Preclinical** evidence.

**Therapeutic hypothesis 2:** Use **PRMT5-pathway inhibition** in splicing-dysregulated NSCLC enriched for RBM10 mutation.
**Rationale and evidence:** A SAM-competitive PRMT5 inhibitor study in NSCLC showed that drug sensitivity associated with spliceosome-related states and with mutations in splicing factors including **RBM10**, supporting PRMT5 inhibition as an indirect strategy against RBM10-driven splicing dysregulation (PMID: 34737197).
**Strength: Preclinical/translational** evidence.
**Clinical translation status:** Active PRMT5-pathway trials include thoracic/NSCLC-eligible studies, although they are generally **MTAP-loss selected**, not RBM10-selected; examples include NSCLC/thoracic studies with **TNG462**, **AMG 193**, and **TNG456** (NCT: NCT06922591, NCT06333951, NCT06810544). These are not evidence of RBM10-specific efficacy, but they are relevant pathway-level opportunities.

#### D. RBM10 mutation as a modifier of EGFR-targeted therapy response

**Therapeutic hypothesis:** RBM10 LoF should be viewed primarily as a **negative response modifier** for EGFR TKI therapy, not a positive predictive biomarker for a specific approved drug.
**Rationale and evidence:** Multiple NSCLC studies linked RBM10 mutation to **shorter PFS/OS** or decreased sensitivity with EGFR-TKIs, especially **osimertinib**. RBM10 mutation was an independent adverse factor for PFS/OS in second-line osimertinib settings and was associated with faster progression in broader NSCLC cohorts (PMID: 38323283, 38496877, 39138107). Additional NSCLC data also note reduced EGFR-TKI sensitivity in RBM10-mutated tumors (PMID: 41934032). In a neoadjuvant osimertinib phase II study, RBM10 was among the common co-alterations, but that trial was not designed to prove RBM10-specific benefit (PMID: 39028931).
**Strength:** Mostly **retrospective/observational clinical** evidence.
**Clinical implication:** For RBM10 N446Kfs*35, EGFR-targeted therapy remains driven by the primary oncogenic driver, but RBM10 LoF may justify **closer monitoring for attenuated benefit or earlier resistance** rather than selection of a specific RBM10-directed approved therapy.

#### E. MET-directed rescue after EGFR-TKI resistance when MET amplification emerges

**Therapeutic hypothesis:** In EGFR/RBM10-mutant LUAD that acquires **MET amplification**, consider **MET inhibitor + EGFR TKI** strategies.
**Rationale and evidence:** A clinical case report of EGFR- and RBM10-mutant lung adenocarcinoma described fatal progression after emergence of **MET amplification**, explicitly arguing for earlier recognition and **timely MET-targeted therapy in combination with EGFR-TKI** (PMID: 41878697). This is not proof that RBM10 predicts MET dependence, but it is a clinically relevant resistance-management lesson in RBM10-mutant disease.
**Strength: Case report/clinical observation**.

#### F. Immunotherapy: possible context-dependent signal, but not established actionability

**Therapeutic hypothesis:** RBM10 LoF may sometimes enrich for **ICI benefit**, especially in **EGFR-wildtype LUAD**, but evidence is not yet sufficient to treat RBM10 as a standalone immunotherapy biomarker.
**Rationale and evidence:** In one LUAD cohort, **EGFR-wildtype LUAD with RBM10 LoF** showed

longer PFS on PD-1 inhibitors than non-LoF RBM10-mutant cases, alongside higher PD-L1/TMB features in EGFR-wildtype disease (PMID: 38912481). Another NSCLC study found co-mutation interactions involving RBM10 that influenced ICI efficacy modeling (PMID: 35042953).
**Strength: Retrospective exploratory** evidence.
**Clinical implication:** RBM10 alone is **not actionable enough** to direct immunotherapy choice, but it may contribute to composite biomarker interpretation.

##### G. Pan-cancer mechanistic analogies: NF-κB and JAK/STAT vulnerabilities support broader RBM10-LoF biology

**Therapeutic hypothesis:** RBM10 loss may create additional therapeutic dependencies such as **NF-κB** or **JAK/STAT** signaling, although these are not yet validated in NSCLC.
**Rationale and evidence:** In thyroid cancer, RBM10 loss promoted metastasis and a CRISPR screen identified **NF-κB effectors** as synthetic lethal dependencies, suggesting a therapeutically targetable state downstream of RBM10 loss (PMID: 39992626). In pancreatic cancer, RBM10 deficiency increased JAK/STAT signaling and immune escape, while the **JAK inhibitor AZD1480** restored NK-cell cytotoxicity (PMID: 40740233). These support the broader concept that RBM10 LoF can generate targetable pathway addictions across tumor types.
**Strength: Preclinical**, pan-cancer supportive evidence.
**Clinical implication:** These findings are useful for hypothesis generation but remain **indirect** for NSCLC.

#### 3. Conclusion:

For **RBM10 N446Kfs*35** in NSCLC, the evidence supports a **LoF tumor-suppressor alteration** with **no established approved RBM10-directed therapy**. The most credible therapeutic leads are **WEE1 inhibition** and **S1P-pathway inhibition/fingolimod** as mechanism-based investigational strategies, with additional support for **RAC1B/splicing-directed approaches** and awareness that RBM10 mutation may **dampen EGFR-TKI efficacy**. Clinically, this variant is best viewed today as a **biologically meaningful, trial-informing alteration** rather than a standard-of-care actionable biomarker.

#### 4. Curated Evidence Lists:

Relevant PMIDs: ["39080280", "40069781", "40548642", "34737197", "38323283", "38496877", "39138107", "41934032", "39028931", "41878697", "38912481", "35042953", "39992626", "40740233"]
Relevant NCTs: ["NCT06424067", "NCT06922591", "NCT06333951", "NCT06810544"]
