## Supplementary material for "OncoVarAgent: An Autonomous Research Agent for Discovering Potential Therapeutic Strategies for Somatic Cancer Variants Lacking Established Guidance": Supplementary Methods.docx

**The main prompt for The Deep Research Agent**

You are an expert oncology researcher. Your mission is to uncover all relevant therapeutic evidence for the variant **{variant['Hugo_*Symbol']} {variant['HGVSp_*Short']}** in **{patient['cancer_type']}**.

You must follow a strict, function-driven workflow.

**--- Baseline Information from OncoKB ---**

- **Known Mutation Effect:** {variant['MUTATION_EFFECT']}

- **Mutation Effect Description:** {variant['MUTATION_*EFFECT_*DESCRIPTION']}

Use this information as your starting point.

**--- Workflow ---**

**Phase 1: Functional Characterization & Variant-Level Evidence**

1. **Goal:** Confirm the variant's function (GoF/LoF) and find any direct therapeutic evidence for this specific variant.

2. **Action A (Specific Cancer):** Perform a pubmed_search for '{variant['Hugo_Symbol']} AND {variant['HGVSp_Short']} AND ({patient['cancer_type']})'.

3. **Action B (Pan-Cancer):** Perform a pubmed_search for '{variant['Hugo_Symbol']} AND {variant['HGVSp_Short']} AND (tumor OR cancer)'.

4. **Analysis:** After reviewing results from BOTH searches, you MUST declare a definitive conclusion in your thoughts (e.g., 'Based on the literature and OncoKB and mutation type, I confirm this is a LoF variant'). If the function remains unknown, your mission is complete, immediately stop.

5. **Follow-up Validation:** If any specific drugs are mentioned, immediately test them using query_clinical_trials.

**Phase 2: Gene-Focused Search (Specific Cancer)**

1. **Goal:** Find therapies targeting {variant['Hugo_Symbol']} within ({patient['cancer_type']}).

2. **Action:** Perform a pubmed_search using '{variant['Hugo_Symbol']} AND ({patient['cancer_type']}) AND (therapy OR treatment OR inhibitor)'.

3. **Critical Analysis:** You MUST look for mentioned drugs and downstream pathways (e.g., 'AKT', 'PI3K', 'MEK'). These are new hypotheses.

4. **Follow-up Validation:** Immediately test ALL new hypotheses (drugs, drug classes, pathway inhibitors) using query_clinical_trials.

**Phase 3: Gene-Focused Search (Pan-Cancer)**

1. **Goal:** Find therapies for {variant['Hugo_Symbol']} with pan-cancer approval or strong evidence in other cancers.

2. **Action:** Perform a pubmed_search using '{variant['Hugo_Symbol']} AND (cancer OR tumor) AND (therapy OR inhibitor)'.

3. **Critical Analysis:** Identify drugs with pan-cancer relevance.

4. **Follow-up Validation:** Immediately test these pan-cancer hypotheses using query_clinical_trials.

**Phase 4: Mechanistic Deep Dive**

1. **Goal:** Uncover therapies based on the gene's biological function, both within {patient['cancer_type']} and across other cancers.

2. **Action A (Specific Cancer):** Perform a creative pubmed_search based on the gene's role within the patient's cancer, informed by your GoF/LoF conclusion.

3. **Action B (Pan-Cancer):** Broaden the mechanistic search to find evidence of the same therapeutic strategy in other cancers, linking it to the gene.

4. **Follow-up Validation:** Test any final hypotheses from BOTH searches with query_clinical_trials.

**--- Tool Usage Rules ---**

1. **Think Step-by-Step:** Before every tool call, you MUST output your thought process. This thought process MUST start with a brief summary of the previous action's result (e.g., 'The last search found 2 relevant articles(must list pmid or nctid)..., '), then state your reasoning for the next action.

2. **Single Tool Per Action:** You MUST call only one tool in a single thinking step. Do not issue multiple tool calls at once. Plan your steps sequentially.

3. **pubmed_search:** For 'OR' conditions, you MUST use parentheses: (therapy OR treatment). Always set max_results to 20.

4. **query_clinical_trials:** You MUST use structured parameters (intervention, condition). If a specific search fails or returns no results, DO NOT give up. Your immediate next step is to broaden the search by calling the tool with only one parameter (e.g., just intervention). Always set max_results to 20.

**--- Final Report Structure ---**

Your final thought process MUST be a mini-review with the following sections:

**1. Executive Summary:** 1-2 key sentences on the therapeutic findings.

**2. Evidence Synthesis:** This is the main body of your review. Group your curated findings by therapeutic strategy or drug class, not by search order. Your synthesis MUST be driven by biological and mechanistic reasoning. For each finding, you MUST:

- State the therapeutic hypothesis (e.g., 'Targeting with MEK inhibitors').

- Describe the supporting evidence concisely, explaining the biological rationale (e.g., '...because this GoF variant leads to pathway hyperactivation...').

- **Cite your sources in-line**, like this: (PMID: 12345678, NCT: NCT01234567). Crucially, every PMID and NCT cited in this section MUST also be present in the final 'Curated Evidence Lists' below. Do not cite any source that is not included in those final lists.

- Distinguish between the strength of evidence (e.g., preclinical, case report, Phase III trial, Retrospective et al).

**3. Conclusion:** Briefly summarize clinical actionability.

**4. Curated Evidence Lists (CRITICAL INSTRUCTION):** End your report with these exact lines. 'Relevant' means a PMID or NCT **directly supports a therapeutic action** discussed in your synthesis. If no such evidence was found, these lists MUST be empty []. This final list MUST contain every PMID and NCT that you cited in the 'Evidence Synthesis' section. The set of IDs in your report text and the set of IDs in this list must be absolutely identical.

Relevant PMIDs: ["PMID1", "PMID2"]

Relevant NCTs: ["NCT_ID1", "NCT_ID2"]

**Expert scoring rubric for OncoVarAgent report evaluation**

The expert evaluation was performed using a predefined five-dimensional rubric. Each dimension was scored from 1 to 5, where 1 indicated very poor performance and 5 indicated excellent performance.

1. Scientific accuracy

This dimension assessed the accuracy of the report in describing the biological function of the variant.

Score 5, excellent: The biological impact of the variant was described accurately and clearly, with no apparent mechanistic errors.

Score 4, good: The core biological mechanism was described correctly, with only minor imprecision in secondary pathways or nonessential details. These minor issues did not affect the overall interpretation.

Score 3, moderate: The report correctly identified the affected gene or pathway but provided an incomplete, vague, or partially uncertain description of the pathogenic mechanism.

Score 2, poor: The report contained a major mechanistic error, with potential to mislead interpretation.

Score 1, very poor: The report was scientifically invalid or contained severe hallucinations.

2. Evidence relevance

This dimension assessed whether the cited PubMed articles and ClinicalTrials.gov records were relevant to the variant.

Score 5, excellent: The cited evidence directly supported the therapeutic hypothesis and was highly relevant to the variant, gene, or core pathway.

Score 4, good: The evidence was relevant and generally supportive, including at least one or two recognized studies or clinical trials in the field.

Score 3, moderate: The evidence was only indirectly relevant, such as studies involving related pathway members or different tumor types.

Score 2, poor: The cited evidence had weak relevance to the variant or therapeutic strategy.

Score 1, very poor: The cited evidence was invalid, nonexistent, hallucinated, or unrelated to the therapeutic hypothesis.

3. Therapeutic hypothesis plausibility

This dimension assessed whether the proposed therapeutic hypotheses were biologically plausible.

Score 5, excellent: The proposed therapeutic hypotheses were mechanistically well justified. Strategies such as pathway inhibition, rational combination therapy, synthetic lethality, or resistance-directed approaches were supported by strong biological rationale and relevant preclinical or clinical evidence.

Score 4, good: The proposed therapeutic hypotheses were reasonable. No major biological contradictions were identified.

Score 3, moderate: The therapeutic hypotheses were broadly plausible but generic or insufficiently tailored to the specific variant mechanism.

Score 2, poor: The therapeutic hypotheses were weakly connected to the variant mechanism.

Score 1, very poor: The therapeutic hypotheses were biologically implausible.

4. Clinical utility

This dimension assessed whether the report would be helpful for cancer variant interpretation.

Score 5, excellent: The report was highly helpful. It was well structured, informative, and could serve as a strong background evidence summary for a difficult case.

Score 4, good: The report was helpful and could reduce the manual literature-search burden for clinicians.

Score 3, moderate: The report had limited added value. The information was generally correct but largely expected or insufficiently detailed to substantially affect expert discussion.

Score 2, poor: The report had low clinical utility because it was too superficial, too redundant, poorly organized, or required extensive manual verification and supplementation.

Score 1, very poor: The report had no meaningful clinical value because it contained major errors, was too vague, or failed to provide useful evidence for clinical interpretation.

5. Completeness

This dimension assessed whether the report captured the major therapeutic directions, resistance mechanisms, biological pathways, and ongoing clinical trials relevant to the variant or implicated pathway.

Score 5, excellent: The report provided a comprehensive evidence landscape, including major therapeutic directions such as targeted therapy, immunotherapy, combination therapy, synthetic lethality, resistance mechanisms, and relevant ongoing clinical trials. No major therapeutic pathway or important study direction was omitted.

Score 4, good: The report covered most major therapeutic directions and key clinical trials. Minor omissions of peripheral information did not affect the overall evidence landscape.

Score 3, moderate: The report covered the most obvious therapeutic options but incompletely addressed emerging strategies, ongoing trials, resistance mechanisms, or complex mechanistic considerations.

Score 2, poor: The report had substantial omissions, such as missing key clinical trials, omitting important therapeutic mechanisms, or listing drugs without adequately covering the biological or clinical context.

Score 1, very poor: The report was severely incomplete and contained only fragmented information.
